# Comparison of Methodologies for Absolute Binding Free Energy Calculations of Ligands to Intrinsically Disordered Proteins

**DOI:** 10.1101/2024.07.19.604182

**Authors:** Michail Papadourakis, Zoe Cournia, Antonia S. J. S. Mey, Julien Michel

## Abstract

Modulating the function of Intrinsically Disordered Proteins (IDPs) with small molecules is of considerable importance given the crucial roles of IDPs in the pathophysiology of numerous diseases. Reported binding affinities for ligands to diverse IDPs vary broadly and little is known about the detailed molecular mechanisms that underpin ligand efficacy. Molecular simulations of IDP-ligand binding mechanisms can help us understand the mode of action of small molecule inhibitors of IDP function, but it is still unclear how binding energies can be modeled rigorously for such a flexible class of proteins. Here we compare alchemical absolute binding free energy calculation (ABFE) and Markov-State Modelling (MSM) protocols to model the binding of the small molecule 10058-F4 to a disordered peptide extracted from a segment of the oncoprotein c-Myc. The ABFE results produce binding energy estimates that are sensitive to the choice of reference structure. In contrast, the MSM results produce more reproducible binding energy estimates consistent with weak mM binding affinities and transient intermolecular contacts reported in the literature.

## Introduction

Intrinsically Disordered Proteins (IDPs) are composed of protein sequences that are unable to fold spontaneously into stable, well-defined globular three-dimensional structures but are dynamically disordered and fluctuate rapidly over an ensemble of conformations.^1–6^ IDPs are highly abundant in nature, amounting to 40% of eukaryotic, 25% of viral and 10% of bacterial proteins.^3,7^ IDPs participate in protein-protein interactions through a coupled-folding upon binding mechanism, which is characterized by high-specificity low-affinity complexes due to the high entropic cost of complex formation.^8,9^ IDPs play vital roles in signal transduction and transcription^10–13^ and they drive the formation of membrane-less organelles, which play a critical role in the spatio-temporal organization of the cell, as they have the ability to undergo liquid-liquid phase separation.^14,15^

Given their abundance and biological importance, there is a need for chemical agents that can control their function. However, until recently IDPs were considered as undruggable since their considerable flexibility is an inherent challenge for typical medicinal chemistry modalities. Additionally, little is known about the molecular driving forces that underpin IDP recognition, and how such principles can inform the design of man-made molecules that can effectively modulate the function of IDPs.^16^ Yet some studies have demonstrated inhibition of the biological functions of IDPs using small molecules.^17^

Reports in the literature show a broad range of binding affinity measurements for small molecule IDP ligands. Fasudil has been shown to attenuate alpha-synuclein aggregation in *in vivo* models of Parkinson’s disease at low doses, although NMR titration experiments suggest Fasudil shows a relatively weak dissociation constant in the 1-3 mM for alpha-synuclein.^18^ Basu *et al.* have also reported a similar disconnect between the ca. 5 mM binding constant of the small molecule EPI-001 for the transactivation domain of the androgen receptor measured by NMR, and a ca. 5 *µ*M IC_50_ activity in a luciferase reporter cell assay.^19^ Heller *et al.* determined, by NMR titration experiments, a *K*_D_ of ca. 6 *µ*M for the small molecule 10074-G5 and disordered monomeric amyloid-*β* (A*β*) peptide^20^ and a *K*_D_ of ca. 300 *µ*M for 5-fluoroindoke to the disordered domain of non-structural protein 5A of the Hepatitis C virus. The disconnect between weak high *µ*M to mid mM binding affinities measured in *in vitro* biophysical assay contexts and low micromolar IC_50_ values observed in cellular assays suggest that the mode of action of these small molecules may be more complicated than reversible stoechiometric non-covalent binding.

Molecular simulations have the potential to provide deep insights into the molecular basis of the mode of action of these compounds that has eluded experimental methods to date. However, simulations of small molecule binding to IDPs present challenges for established methodologies. Alchemical free energy methods have become established for the computation of binding affinities of small molecules to well-folded globular proteins.^21,22^ Yet it is unclear whether existing AFE protocols are applicable to ‘fuzzy’ IDP:small molecule complexes.^23^

We focus our attention on the oncoprotein c-Myc, a transcription factor frequently over-expressed in many cancers; inhibition of c-Myc function is widely regarded as a ‘holy grail’ of cancer therapies.^24–27^ A possible inhibition strategy is to prevent heterodimerisation of the c-Myc basic-Helix-Loop-Helix-Leucine zipper (bHLHZip) domain with partner protein Max, a step essential for c-Myc to act as transcription factor.^28^ Several small molecules have been reported to achieve this outcome by binding to the disordered monomeric c-Myc bHLHZip domain in a manner that prevents the formation of the ordered Myc/Max heterodimer.^29,30^ In particular the rhodanine-based compound 10058-F4 inhibited the proliferation of HL60 cells with an IC_50_ value of ca. 40 *µ*M Subsequent fluorescent polarization assays indicated that 10058-F4 binds to c-Myc with a *K*_D_ of ca. 5 *µ*M. Additional surface plasma resonance (SPR) experiments by Müller *et al.* determined the direct binding of 10058-F4 with *K*_D_ values of 40±8 *µ*M.^31^ Finally, Heller *et al.* examined the potency of 10058-F4 with Isothermal Titration Calorimetry (ITC) and with van’t Hoff analysis using fluorescence titration experiments at different temperatures. They did not observe any binding with ITC at room temperature because of a low heat of binding, but a binding free energy of -27.6 ± 8.5 kJ/mol at 25 C*^◦^* was determined via a van’t Hoff analysis. This study concluded that entropic contributions are a key factor for the binding of 10058-F4 to the oncoprotein c-Myc.^32^

Careful biophysical studies delineated the binding site of 10058-F4 to c-Myc_402_*_−_*_412_, and a model of 10058-F4 bound to c-Myc_402_*_−_*_412_ was proposed on the basis of NMR experiments. This region is located at the interface between the H2 and Zip region in the c-Myc-Max dimer and forms a hydrophobic cluster of Tyr402, Ile403, Leu404, Val406, Ala408 in the c-Myc_402_*_−_*_412_-10058-F4 complex.^30^ Intrigued by this opportunity, Michel and Cuchillo employed molecular dynamics and bias-exchange metadynamics simulations to provide new insights into the mechanisms of molecular recognition between the small molecule 10058-F4 and c-Myc. They reported that the ligand does not have a dominant binding mode but interacts with multiple binding sites through weak and non-specific interactions. Moreover, the compound made preferential contacts with the most hydrophobic region of this sequence, a result that was in agreement with Hammoudeh *et al*.^30,33^ Therefore, this study has highlighted the lack of specificity between 10058-F4 and its target as well as the difficulty of locating possible binding sites for c-Myc. These findings were later replicated by other groups for 10058-F4 and other related ligands.^32,34^

To date, however, MD studies of c-Myc binders or other IDPs have focused on qualitative description of the mechanisms of small molecule-IDP recognition. As a consequence, while this system has been the subject of simulation studies by several groups, no simulated binding affinities have been reported.^32,34^ In this study, we benchmark two protocols to estimate the standard binding free energy of a small molecule to an IDP. We first use an alchemical free energy calculation protocol to compute the binding free energy of the small molecule 10058-F4 to a disordered peptide from the protein c-Myc. We compare the computed ensembles and binding energy estimates from the absolute binding free energy methods (ABFE) with similar quantities obtained by Markov State Modelling protocols. MSMs lend themselves to this problem, as they provide both kinetic, as well as populations, such as bound and unbound states, from simulations in a mathematically rigorous fashion.^35,36^ Overall, our results inform the applicability of these two molecular simulation methods to the study of small molecules:IDP interactions.

## Methods

### MD simulations of c-Myc/ligand complexes

For the MD simulations, NMR constraints as reported by Hammoudeh *et al.* were used to construct the initial binding pose for the c-Myc_402_*_−_*_412_/10058-F4 complex.^30^ This binding pose served as the input structure for the initial alchemical free energy calculations (Hammoudeh pose and Heller’s parameters) as well as for the two sets of equilibrium MD simulations.

Two sets of equilibrium MD simulations were performed for this study. The first set was implemented to derive the most stable pose as an initial conformation for the absoluteFEP protocol for the protein-ligand complex. For this purpose, the input files used for these simulations were created using the software FESetup1.2.1.^37^ Proteins were parameterized using ff14SB Amber force field,^38^ while GAFF2 parameters,^39,40^ that use AM1-BCC charges^41^ were assigned to the ligands. The systems were solvated in a rectangular box with TIP3P waters^42^ with a minimum distance between the solute and the box of 12 Å. Counter-ions were also added to neutralize the total net charge.

For the equilibration protocol, energy minimization of the entire system was implemented with 1000 steps of steepest gradients, using sander. Then, an NVT protocol for 200 ps was performed at 298 K, followed by an NPT equilibration for a further 200 ps at 1 atm. Eventually, a 2 ns MD simulation in an NPT ensemble was run with sander to reach a final density of 1 g cm^−3^. This was followed by a production run simulations of the protein-ligand complex for 500 ns using SOMD1 software (revision 2019.1.0) in an NPT ensemble.^43,44^ Temperature control was maintained by an Andersen thermostat with a coupling constant of 10 ps^−1^. Pressure control was achieved with a Monte Carlo barostat that attempted to scale the isotropic box edge every 25 fs. A 10 Å atom-based cutoff distance for the non-bonded interactions was used, using a Barker-Watts reaction field, with a dielectric constant of 78.3. The final coordinate files were retrieved with cpptraj.

The second set of MD simulations includes three individual simulations run for 20 *µ*s each. These simulations are divided into 20 replicas of 100 ns each, which will be used to construct a Markov state model (MSM) of the c-Myc_402_*_−_*_412_/10058-F4 complex. Each set of simulations was performed with a different force field, two specifically designed for IDPs and the third a standard amber protein force field (ff14SB). The first set of input files for the protein-ligand complex was generated with the same method and force fields (ff14SB Amber force field for the protein and GAFF2 parameters for the ligand) as for the first set of MD simulations. The same equilibration protocol was used, and the final coordinate file was obtained with cpptraj. The protein-ligand complex was run using 20 replicas of 100 ns each using the SOMD1 software (revision 2019.1.0) in the NPT ensemble at 300 K and 1 atm. A 2 fs timestep was used, and all bonds involving hydrogens were constrained. The temperature control was maintained by an Andersen thermostat with a coupling constant of 10 ps^−1^. Pressure control was achieved using a Monte Carlo barostat. Periodic boundary conditions were used with a 10 Å atom-based cutoff distance for the non-bonded interactions together with a Barker Watts reaction field with a dielectric constant of 78.3 for the electrostatic interactions.

For the second simulation the ff14IDPSFF Amber force field^45^ was selected for c-Myc_402_*_−_*_412_ as it is a specific force field for IDPs, GAFF2 parameters,^39,40^ with AM1-BCC partial charges for the ligand through the LEaP module in the Amber 17 suite.^46^ The model was then solvated in a rectangular box of TIP3P water molecules and charge neutrality was enforced through the addition of the necessary counter ions. The input coordinates were energy minimized using 5000 steps of steepest gradients with heavy protein atoms that were position restrained with a force constant of 1000 kJ mol^−1^ nm^−2^. The system was then equilibrated for 100 ps using an NVT ensemble and the same restraints as in the previous step. Finally, 100 ps of NPT ensemble at 1 atm were performed to reach a final density of about 1 g cm^−3^. Next, GROMACS 5.0.5^47^ was used to perform 20 replicas of 100 ns each for the protein-ligand complex in the NPT ensemble at 300 K and 1 atm. A 2 fs timestep was used and LINCS^48^ algorithm was employed to constrain bonds involving hydrogen. Temperature control was maintained at 300 K with a stochastic Berendsen thermostat,^49^ and pressure was achieved using a Parrinello-Rahman barostat.^50^ Electrostatic interactions were handled using Particle Mesh Ewald with a Fourier grid spacing of 1.6 Å. Van der Waals interactions were handled using the Lennard-Jones potential.^51^ The cut-off distance for non-bonded interactions was set at 12 Å, with a switching function applied beyond 10 Å.

The third force field tested was Charmm36m which has been parameterized for IDPs,^52^ GROMACS 5.0.5 package^47^ was used to prepare the third set of input files. The general Charmm force field^53^ was selected for the ligand. The model was then solvated in a rectangular box with TIP3P waters^42^ with a box length of 12 Å away from the edge of the solute. In addition, counterions were added to neutralize the total net charge. A similar equilibration and production protocol as for the previous setup was followed to produce 20 *µ*s MD simulations.

### Double decoupling protocol

Alchemical free energy simulations were performed using a standard double decoupling protocol implemented in the SOMD1 (Figure S1).^54^ Detailed protocols are described in the Supporting Information (SI).

Free energy changes were estimated with the multistate Bennet acceptance ratio method as implemented in the Sire utility *analysefreenrg*.^55^ To achieve a more robust estimation of free energies, each simulation was repeated three times, using different initial velocities drawn from the Maxwell-Boltzmann distribution and statistical uncertainties are reported as one standard error of the mean.

### Markov State Modelling protocol

The resulting pool of trajectories from the s long set of MD simulations was used to construct MSMs for the three different force fields using the pyEMMA 2.3.0 software package. ^56^ Three different features, chosen based on chemical intuition and represent the distance between the protein and the ligand, were used to cluster the MD simulations and construct MSM models. All features involved distances between the selected atoms of 10058-F4 and the C*_a_* atoms of the c-Myc peptide (Figure S2).

The first metric required the calculation of eleven distances between the nitrogen atom of the ligand and the CAs of each amino acid of the c-Myc peptide in each snapshot (Metric 1). Then, dimensionality reduction was performed using tICA to construct a low dimensionalrepresentation of the data.^57^ However, tICA retained ten out of eleven dimensions to explain 95 % of the slow timescales of the system. As a result, we decided to use the original eleven dimensions for each snapshot.

The second metric used only the shortest distance between the nitrogen of the ligand and the CAs of each amino acid of the oncoprotein for every snapshot (Metric 2). The third metric used the shortest distance between either the nitrogen or the carbon atoms highlighted in Figure S2 and the CAs of each residue at each snapshot (Metric 3). For the last two metrics, we did not perform a reduction of the dimensional space as the initial feature space consisted of only one dimension.

Subsequently, k-means clustering using 75 clusters was performed to discretize the trajectories and obtain microstates for the MSM construction. Implied timescales (ITS) of the dominant eigenvectors were calculated for each metric (Figures S3-S5) to identify the optimal lag time. This, along with the default parameters of pyEMMA, was used to estimate the MSM transition matrices using the Bayesian MSM. The validity of the MSMs was tested with the Chapman-Kolmogorov test (CK test) where the full transition probability matrix T was coarse-grained into 2 metastable states (Figures S6-S8).^58^ Then, we performed spectral clustering using PCCA++ algorithm to coarse-grain the microstates into two metastable states.^59^ In addition, the stationary probabilities (*π*) of the two metastable states were calculated by summing over the populations of the 75 microstates. The Mean First Passage Times (MFPT) between the two states were estimated from the Bayesian MSM.

The standard binding free energy of the ligand was estimated using equation 1.

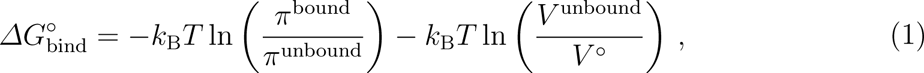

where *k*_B_ is the Boltzmann constant, *T* is the temperature in Kelvin, *π* accounts for the stationary probabilities of the bound and unbound macrostates. The second term corrects for the volume of the unbound state in the simulation box is different from the standard volume conditions for a 1 M dilute solute (*V ^◦^* = 1660 Å^3^/mol).

To determine *V*_bound_, the volume of space available to the ligand in the unbound state, we computed the average distance between the center of masses of the ligand and the protein from 1000 snapshots sampled from the bound macrostate. We then estimated the bound volume *V*_bound_ as the volume of a sphere with a radius equal to this average distance. We also used cpptraj to compute the average volume *V_total_* of the simulation box from these 1000 snapshots. The volume of the unbound state was then taken as the difference between *V*_total_ and *V*_bound_.

Finally, we also computed the rate constants *k*_on_ and *k*_off_ for the binding process by using the calculated MFPTs between bound and unbound states. Assuming first-order reactions, the relation between the rates and the corresponding MFPTs is provided by the following equations:^60^

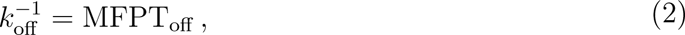

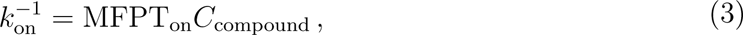

where C_compound_ is the concentration of the ligand in the simulation box.

## Results and discussion

### Binding free energies from the alchemical ABFE protocol

A double decoupling absoluteFEP protocol was employed to reproduce the binding affinity and binding site preference of the known c-Myc binder 10058-F4 to the 402-412 Myc fragment.^23,61^ The starting points for the MD simulations and the absoluteFEP protocol of the c-Myc_402_*_−_*_412_/10058-F4 complex are shown in Figure 1 along with the binding free energies computed from the absoluteFEP protocol. A breakdown of the components obtained along the thermodynamic cycle is shown in the Supporting Information (Table S1).

**Figure 1:**
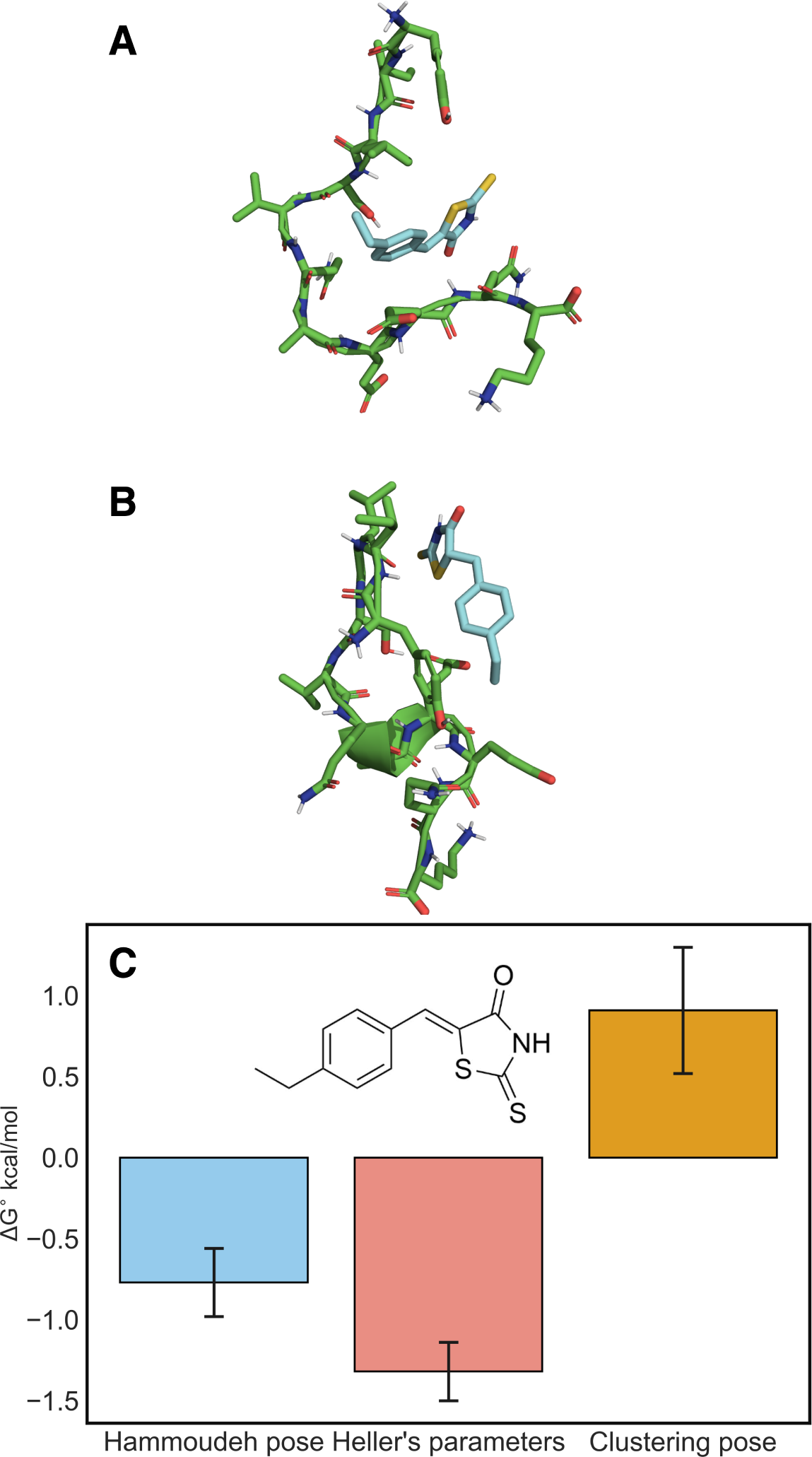
A) Starting structure of c-Myc_402−412_/10058-F4 complex for the absoluteFEP protocol (Hammoudeh pose).^30^ B) Starting structure taken from the most stable cluster in the 500 ns c-Myc_402−412_/10058-F4 equilibrium simulation for absoluteFEP protocol (Clustering pose). C) Computed standard free energies of binding for 10058-F4 in complex with c-Myc_402−412_ using the absoluteFEP protocol.

The protocol yielded reproducible results between the three independent runs, but suggests only a very weak affinity for 10058-F4 (−0.8 ± 0.2 kcal/mol) which contrasts with the previously reported experimental results. A possible reason for this inconsistency could be the force field parameters of 10058-F4. In 2017, Heller *et al.* reported a custom parameterization of the ligand as they observed that GAFF poorly represented the torsional energetics of this ligand.^32^ Thus, we repeated the absoluteFEP protocol using the customized parameters for 10058-F4. However, the resulting binding free energy of the molecule was only slightly more negative (−1.3 ± 0.2 kcal/mol).

One possible reason for the poor computed energetics could be that the conformation of the c-Myc peptide was not representative of the dominant binding mode observed experimentally. To test for this we carried out 500-ns long MD simulations of the c-Myc/10058-F4 complex. Subsequently, we performed clustering with the k-means algorithm and the RMSD of the ligand versus the protein as a metric using cpptraj. The most representative binding pose for the complex (Figure 1C) was then selected and used as the initial conformation for the absoluteFEP protocol.

The binding free energies computed using those structures were even more positive than seen previously for 10058-F4 (0.9±0.4 kcal/mol). Furthermore, there was limited evidence of a dominant binding mode in the MD simulation, with the compounds reversibly binding and unbinding several times. This suggested challenges for the ABFE protocol that provides limited sampling per window. We, therefore, investigated a second approach to capture the free energy of binding using MSMs.

### Binding free energies from Markov State Modelling protocols

Extensive 20 ms-long MD simulations of the c-Myc_402−412_/10058-F4 complex were conducted using three different force field parameter sets. The rationale behind these simulations was to examine the behavior of the c-Myc-ligand complex when protein force fields developed specifically for IDPs are used. Three different metrics, which are dependent on the distance between the protein and the ligand, were used to construct three MSM models for each force field from the simulation data using pyEMMA software. The ten slowest ITS for each metric and each force field were plotted for a range of lag times, *τ*. The corresponding plots are provided in the SI (Figures S3-S5).

The lag time chosen for all analyses was 200 ps. The Markovianity of the different models was validated through CK tests. The resulting plots are given in the SI (Figures S6-S8). Small deviations are observed in the CK test, but deemed adequate enough to proceed with the analysis at a lagtime *τ* of 200 ps.

The CK tests for the second metric show a small deviation from the kinetic behavior of the system on longer timescales. Thus, a lag time of 200 ps is an appropriate choice to predict the long-timescale behavior of the three systems. After successful statistical validation of the models, a two-macrostate MSM model was constructed for each simulation using PCCA++. The grouping turned out to distinguish different protein-ligand conformational states and separate them as bound and unbound based on the distance between the ligand central atom, and the CA atoms of the c-Myc peptide. The probability distribution of distances for the three metrics in bound and unbound states are depicted in the SI (Figures S9-S11). Overall there is a clear preference for microstates in the ‘bound’ macrostate to show lower distances between the ligand and the peptide than in the ‘unbound’ macrostate, however, some overlap remains, which could be due to the broad range of MD snapshots assigned to a single microstate.

The stationary probabilities of these states, *π*_1_ for the unbound state and *π*_2_ for the bound state, were computed by summing over all the microstates and are reported in Table S2. The values of the stationary distributions highlighted that the bound state was more dominant for metrics 2 and 3 but it differed in population amongst the three force fields. On the other hand, metric 1 showed that the unbound state was the most favorable for ff14SB and Charmm36m. It was apparent that the Amber IDP force field showed the strongest tendency for 10058-F4 to bind favourably to the c-Myc_402−412_ peptide.

To provide a more quantitative interpretation of this binding process, we calculated both the binding affinity and the kinetics of the overall process of binding for the three metrics for each force field. The overall binding free energy was computed from the stationary distributions and the standard state correction term in equation 1. The values obtained from this calculation for the first two metrics were similar between the three force fields and were on average 1.5 kcal/mol more negative than the most negative binding energies determined with the absoluteFEP protocol (Figure 2). However, for the case of metric 3, the binding free energies computed for Charmm and FF14SB were similar to the binding free energy obtained from the absoluteFEP protocol, when custom parameterisation was used, and differed from the corresponding binding free energy of the Amberidp force field. The most negative binding energy estimates (-2.7 kcal/mol) are obtained with Amberidp/GAFF2, which corresponds to a dissociation constant of approximately 10 mM. Such affinity estimates are in the range of NMR measurements used to characterize the binding of small molecules to the IDPs *α*-synuclein (1-3 mM) and the transactivation domain of the androgen receptor (5 mM).^18,19^ They do however deviate more substantially from K_D_ values derived from fluorescence polarisation measurements reported for 10058-F4.

**Figure 2:**
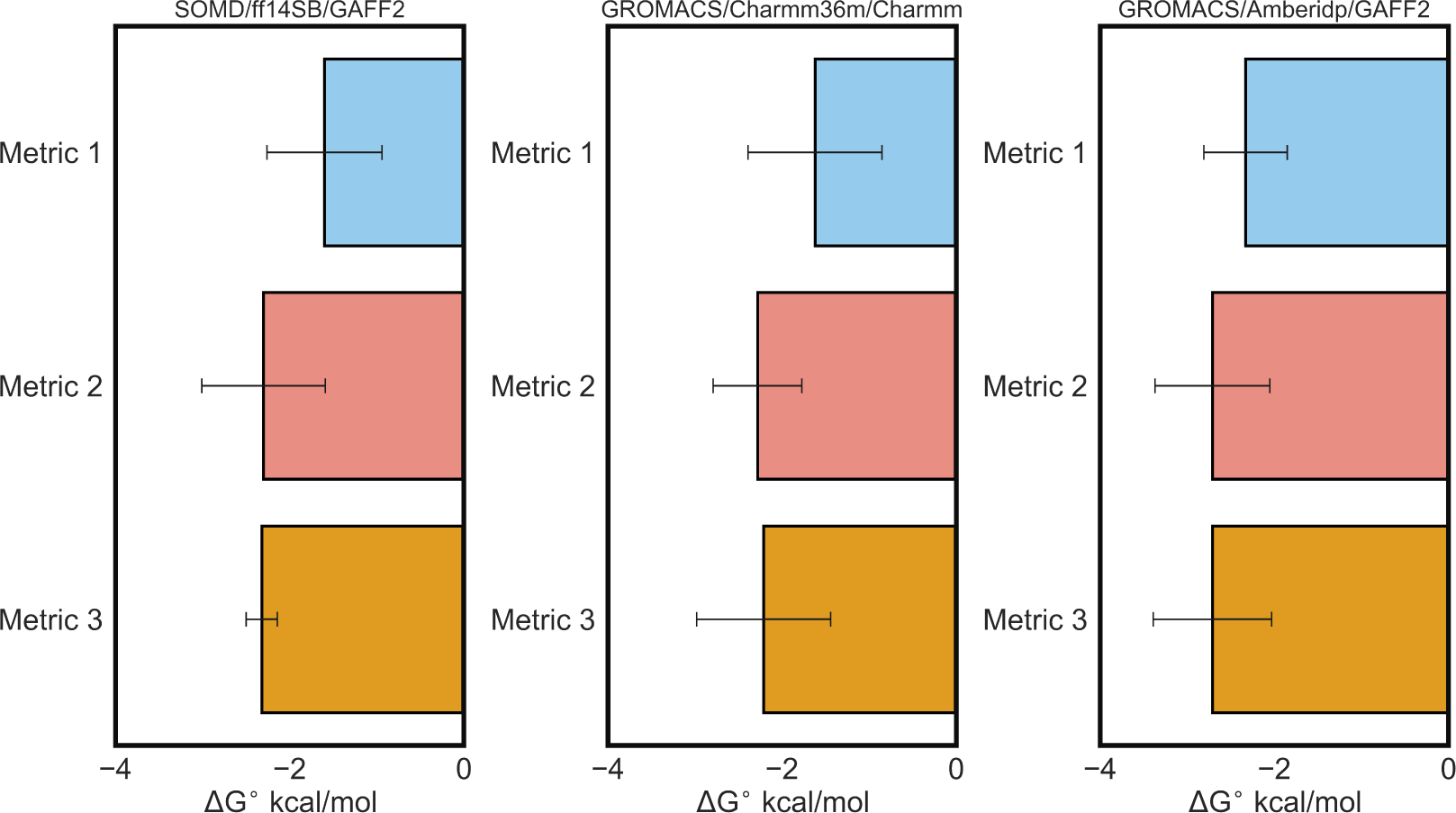
Computed standard free energies of binding for 10058-F4 in complex with c-Myc_402−412_ using the MSM protocol.

Another important feature that can be computed from the MSM models is the kinetics that govern the binding process of 10058-F4 to c-Myc. For this purpose, Mean First Passage Time values (MFPTs) between the two states in each force field for the three metrics were calculated from the Bayesian MSM. MFPT values were then converted into rate constants for the binding and unbinding of 10058-F4 to c-Myc (using a concentration of the ligand equal to 0.02 M given the box dimensions). The k*_on_* and the k*_off_* values for each of the three force fields for the three metrics are illustrated in Figure 3 and summarized in Table S4.

**Figure 3:**
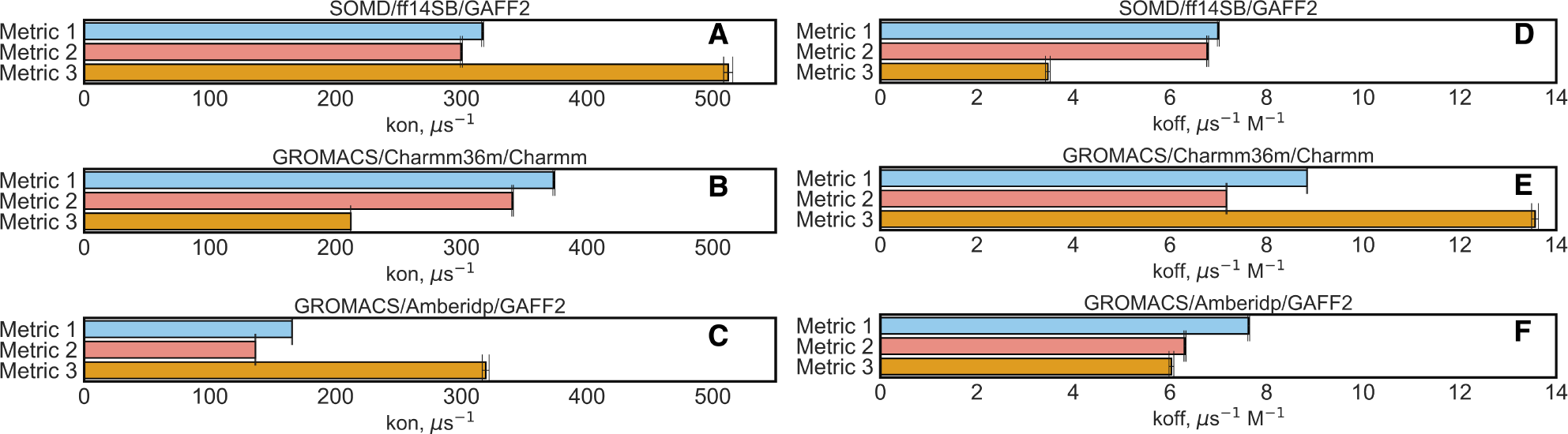
The kinetic reaction rates (k_on_ and k_off_) of the three different force fields for the three metrics for the bound and the unbound states respectively.

The computed k_on_ and k_off_ values from our MSM models can be compared with values that would be expected for a protein-ligand binding process. The typical range of the k_on_ rates spans between 10^3^ s^−1^ M^−1^ to 10^9^ s^−1^ M^−1^, with the latter corresponding to the rate limit of diffusion of a solute to the solvent.^62^ Thus, the on rate constants computed (ca. 10^8^ s^−1^ M^−1^) from the MSMs are close to diffusion limit. k_off_ values typically range from 1 s^−1^ to around 10^7^ s^−1^, due to the long-lasting nature of protein-ligand interactions.^62^ The MSM-derived k_off_ values appear therefore to be at the upper range of what is experimentally observed. Overall, the picture that emerges is one of weak affinity and very fast binding/unbinding kinetics.

The final aim of this study was to identify the residues that the ligand prefers to interact with when in its bound state. For this purpose, 1000 snapshots were extracted based on microstate probabilities using the appropriate pyEMMA functions in order to create a trajectory with snapshots from the bound state of metric 2. This procedure was repeated three times. Metric 2 showed the strongest tendency for 10058-F4 to bind favourably to the c-Myc_402−412_ peptide for all applied force fields. In the resulting trajectories, we applied a custom Python script using the *mdtraj* module to count the number of carbon atoms of the ligand and measure the distance of those atoms with the carbon atoms of the protein. This metric allowed us to evaluate the hydrophobic contacts of the ligand with every residue for each snapshot. A cutoff of 4 Å was used for every distance to only identify close contacts between 10058-F4 and each residue of the oncoprotein c-Myc. We also examined the ability of the ligand to engage in hydrogen bonding interactions with each residue using the cpptraj module. The total number of hydrophobic contacts and hydrogen bonds formed during the 1000 snapshots for each of the 4 trajectories for the Amberidp force field are depicted in Figure 4A and 4B respectively, while the total number of contacts for the other two force fields are given in the SI (Figures S10 and S11).

**Figure 4:**
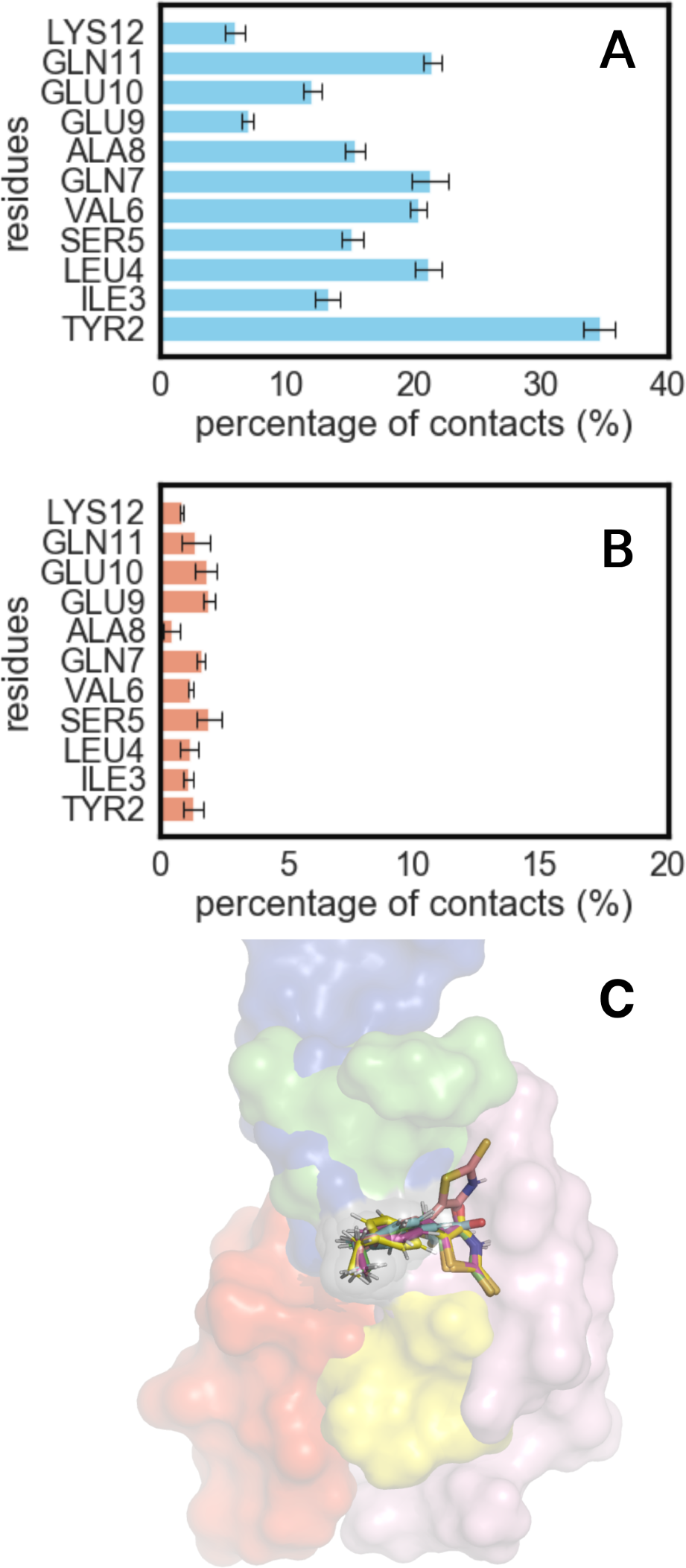
A) Mean percentage of hydrophobic contacts formed during the four trajectories, each containing 1000 snapshots and B) Mean percentage of hydrogen bonds formed during the four trajectories, each containing 1000 snapshots for Amberidp force field. C) Five representative snapshots from the bound state of metric 2 for the AmberIDP force field, each depicting a distinct conformational state of the protein in a different color.

The results for each force field indicated that the ligand prefers to bind in the N terminus of c-Myc_402−412_, especially with Tyr402. In addition, the binding process is mainly characterized by van der Waals interactions rather than hydrogen bonds, since only a small fraction of the 1000 snapshots involve hydrogen bonding interactions between the ligand and the peptide. Five representative snapshots for the Amberidp-Metric 2 analysis were obtained from the corresponding trajectory and are depicted in Figure 4C.

These snapshots highlight the flexibility of the bound state, as c-Myc and 10058-F4 adopt diverse conformations. Finally, we observe that the nature of the bound state described by the MSM appears to be in line with the findings from the previous metadynamics study of Michel and Cuchillo.^33^

## Conclusions

Two molecular dynamics simulation protocols were established to study the interactions of small molecules with the intrinsically disordered protein c-Myc. Alchemical free energy calculations were first applied to compute the absolute binding free energy of the c-Myc_402−412_/10058-F4 complex. This protocol generated reproducible results for this system, but the computed free energies of binding (ca. +1/-1 kcal.mol^−1^) deviated significantly from experimental data (ca. -6 kcal.mol^−1^).

Next, extensive MD simulations were carried out to build Markov state models describing reversible binding/unbinding of 10058-F4 to c-Myc_402−412_. The binding/unbinding kinetics of the protein-ligand complex can be described as a two-state process because the slowest transitions are due to the binding process. We discretized the MD trajectories into bound and unbound states by using as features the distances between some parts of the ligands and the CA atoms of the c-Myc residues.

The binding free energies obtained from these models were more negative than those obtained by the ABFE protocol and also more consistent across different choices of forcefields. The computed conformational ensembles are consistent with an earlier study from the group using a bias-exchange variant of the metadynamics method (BEMD) to extensively sample the energy landscape of c-Myc_402−412_/10058-F4 complex.^33^ The current study indicates that protein amino acids located in the N-terminus of c-Myc_402−412_ were more crucial for binding and the interactions with the ligand were mainly hydrophobic in nature, with similar results found previously. However, the calculated binding affinity is consistent with weak mM binding. This contrasts with the low *µ*M K_D_ values reported for this system in the literature,^30^ but is more consistent with binding constants measured for other IDPs:small molecule interactions.^18,19^ A possible reason for this inconsistency could be that the mechanism of binding of the ligand to c-Myc is more complex than the 1:1 stoichiometry assumed by the molecular models used in this study.

Overall, this study suggests that MSM protocols may have an advantage over ABFE protocols to characterize the binding energies of ligands to IDPs owing to the ‘fuzziness’ of the bound state. The use of protein-ligand restraints implicit in ABFE calculations to define a bound state is problematic when the bound state is extremely conformationally flexible. Additional simulation studies of small molecule IDP interactions are warranted to investigate more complex mode-of-actions such as non-stoichiometric binding, or covalent modifications.^63^

## Notes

The authors declare the following competing financial interest(s): J.M. is a member of the Scientific Advisory Board of Cresset.

### Data and Software Availability

The input files generated during this study are available on GitHub at https://github.com/michellab/idpabfe

## Acknowledgement

M.P. thanks the Principal’s Career Development Scholarship Scheme provided by the University of Edinburgh for PhD studentship funding. M.P. has also received support from the Stamatis G. Mantzavinos’ Postdoctoral Research Scholarships Programme by the Bodossaki Foundation and the HPC-Europa3 Scholarship. Computational time granted from the Greek Research Technology Network (GRNET) in the National HPC facility - ARIS under project ID hpce3007 is acknowledged. The authors thank Dr. Heller for providing the small molecule parameters for 10058-F4.

## Supporting Information Available

The Supporting Information is available free of charge. Additional information and supporting figures and tables describing the alchemical binding free energy calculations’ theory and results concerning different absoluteFEP protocol simulations, stationary probabilities, the average distance between the com of the ligand and the com of the protein in the bound macrostate, volumes of the bound and unbound states, standard binding free energies, Mean first passage times (MFTPs), kinetic reaction rates results from the MSM protocol, metrics used for the MSM protocol, Implied time scales plots and Chapman-Kolmogorov test plots and the total number of contacts from the MSM protocol. (PDF) Scripts and datasets to reproduce the main findings from this study are available at https://github.com/michellab/idpabfe.

## Author Information

### Authors

**Michail Papadourakis** - *EaStCHEM School of Chemistry, University of Edinburgh, David Brewster Road, Edinburgh EH9 3FJ, United Kingdom*; *Biomedical Research Foundation, Academy of Athens, 4 Soranou Ephessiou, 11527 Athens, Greece*; Orcid https://orcid.org/0000-0002-9969-5871;

**Zoe Cournia** - *Biomedical Research Foundation, Academy of Athens, 4 Soranou Ephessiou, 11527 Athens, Greece*; Orcid https://orcid.org/0000-0001-9287-364X;

### Present Addresses

**Michail Papadourakis** - *Department of Nursing, Faculty of Health Sciences, Hellenic Mediterranean University, Heraklion, 71004 Crete, Greece*;

