## Supplementary material for "Comparison of Methodologies for Absolute Binding Free Energy Calculations of Ligands to Intrinsically Disordered Proteins"

### Methods

To calculate the binding free energy of a protein-ligand system PL, the ligand L is annihilated both in the solvated and the protein-bound phase by first switching off the charges and then the van der Waals parameters. An example of the thermodynamic cycle used in these calculations is illustrated in Figure S1. During the complex simulations, a flat-bottom distance restraint protocol is applied to prevent the ligand from drifting away from the binding site. A standard state correction term is applied to relate the volume available to the restrained but non-interacting ligand to standard state conditions. Details of the restraint protocol used for these simulations are fully described in Papadourakis *et al.*[1]

The overall cycle gives a standard free-energy of binding as shown by eq 1:

$$\Delta G^{\circ}_{\text{bind}} = \left(\Delta G^{\text{solv}}_{\text{elec}} + \ \Delta G^{\text{solv}}_{\text{vdW}}\right) - \ \left(\Delta G^{\text{host}}_{\text{elec}} + \ \Delta G^{\text{host}}_{\text{vdW}}\right) + \Delta G^{\circ}_{\text{restr}} \ . \tag{1}$$

For the complex and solvated phases the *discharging* steps were run with nine equidistant $\lambda$ windows and 16 $\lambda$ windows (0.00, 0.05, 0.10, 0.15, 0.20, 0.25, 0.30, 0.35, 0.40, 0.45, 0.50, 0.55, 0.60, 0.70, 0.85, 1.00) were employed for the *vanishing* step, both in bound and free phase.

Each lambda value was simulated for a duration of 10 ns with SOMD in the NPT ensemble. Temperature control was achieved with an Andersen Thermostat with a coupling constant of 10 $ps^{-1}$.[2] Pressure control was maintained by a Monte Carlo barostat that attempted isotropic box edge scaling every 100 fs. A 12 Å atom-based cutoff distance for the nonbonded interactions was used, using a Barker-Watts reaction field with a dielectric constant of 78.3.[3] In the bound phase the restraints parameters of eq. 3.1 were: $R_{ji}$ = 7 Å, $D_{ji}$ = 2 Å and $k_{ji}$= 10 kcal $mol^{-1}$ $Å^{-2}$. The alpha carbons ($C_a$) of residues $Leu_{404}$ and $Gln_{410}$ were chosen as the restraint set of the host atom, while the central carbon atom of 10058-F4 was the corresponding guest atoms.

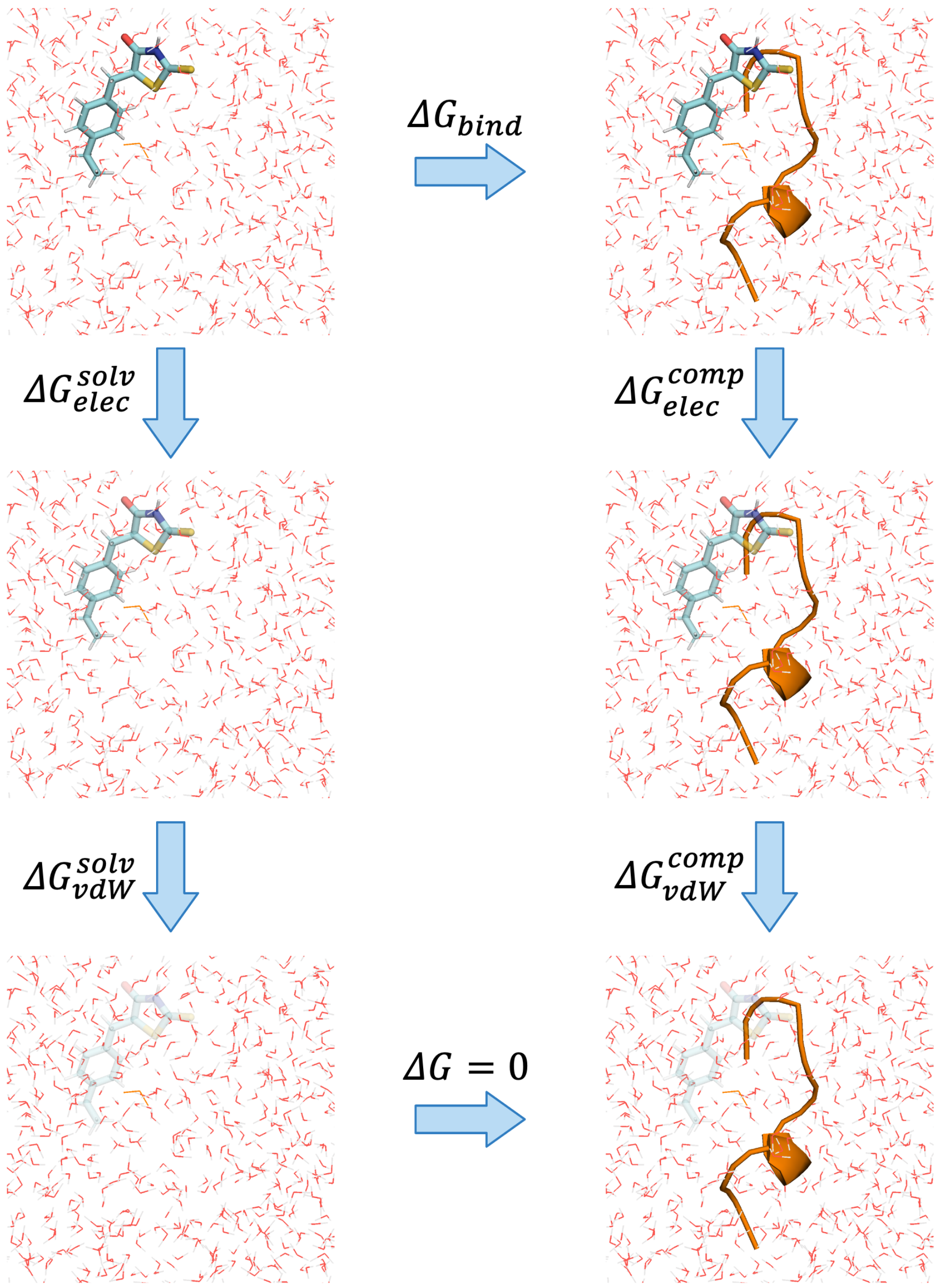


Figure S1: An example of the thermodynamic cycle for standard binding free energy calculations. First, the fully coupled 10058-F4 ligand is simulated in both solvent (top left) and complex phase (top right). Then the charges and the van der Waals interactions are switched off, resulting in a non-interacting 10058-F4 in water (bottom left), and bound to the oncoprotein c-Myc (bottom right).

### Results

Table S1: Results from all the different absoluteFEP protocol simulations for 10058-F4. Energies are reported in kcal/mol

| AbsoluteFEP protocol for 10058-F4 | | | | | | |
|---|---|---|---|---|---|---|
| c-Myc | $\Delta G_{\text{bound discharge}}$ | $\Delta G_{\text{bound vanish}}$ | $\Delta G_{\text{free discharge}}$ | $\Delta G_{\text{free vanish}}$ | $\Delta G_{\text{restraint}}$ | $\Delta G^{\circ}_{\text{ModelC}}$ |
| Hammoudeh pose | 71.67 ± 0.12 | -4.06 ± 0.08 | 72.09 ± 0.00 | -6.76 ± 0.01 | -1.51 ± 0.26 | -0.77 ± 0.21 |
| Heller's parameters | 18.99 ± 0.08 | -4.21 ± 0.16 | 19.25 ± 0.00 | -6.87 ± 0.07 | -1.08 ± 0.12 | -1.32 ± 0.18 |
| Clustering pose | 76.57 ± 0.07 | -5.36 ± 0.36 | 77.47 ± 0.01 | -6.56 ± 0.02 | -1.21 ± 0.05 | 0.91 ± 0.39 |

Table S2: Stationary probabilities of the bound ($\pi_2$) and unbound ($\pi_1$) states for the three metrics of each force-field.

| SOMD/ff14SB/GAFF2 | | |
|---|---|---|
| Metrics | $\pi_1$ | $\pi_2$ |
| Metric 1 | 0.75 ±0.05 | 0.25 ±0.06 |
| Metric 2 | 0.48 ±0.05 | 0.52 ±0.06 |
| Metric 3 | 0.47 ±0.06 | 0.53 ±0.09 |
| GROMACS/Charmm36m/Charmm | | |
| Metric 1 | 0.72 ±0.02 | 0.28 ±0.04 |
| Metric 2 | 0.47 ±0.05 | 0.53 ±0.03 |
| Metric 3 | 0.51 ±0.02 | 0.49 ±0.04 |
| GROMACS/Amberidp/GAFF2 | | |
| Metric 1 | 0.47 ±0.02 | 0.52 ±0.05 |
| Metric 2 | 0.32 ±0.02 | 0.68 ±0.08 |
| Metric 3 | 0.32 ±0.03 | 0.68 ±0.01 |

Table S3: Average distance between the com of the ligand and the com of the protein in the bound macrostate are given in Å, volumes of the bound and unbound states are given in $Å^3$, standard binding free energies are given in kcal/mol.

| SOMD/ff14SB/GAFF2 | | | | |
|---|---|---|---|---|
| Metrics | Average distance | $V_{bound}$ | $V_{unbound}$ | $\Delta G^{\circ}_{msm}$ |
| Metric 1 | 10 ± 4 | 4100 ± 1300 | 75000 ± 1300 | -1.60 ± 0.66 |
| Metric 2 | 10 ± 5 | 5000 ± 2000 | 74100 ± 2000 | -2.30 ± 0.71 |
| Metric 3 | 11 ± 5 | 5000 ± 1700 | 74100 ± 1700 | -2.32 ± 0.68 |
| GROMACS/Charmm36m/Charmm | | | | |
| Metric 1 | 8 ± 2 | 2600 ± 540 | 76000 ± 600 | -1.62 ± 0.77 |
| Metric 2 | 12 ± 5 | 8100 ± 540 | 70000 ± 2700 | -2.28 ± 0.51 |
| Metric 3 | 12 ± 5 | 7700 ± 2500 | 71000 ± 2500 | -2.21 ± 0.77 |
| GROMACS/Amberidp/GAFF2 | | | | |
| Metric 1 | 8 ± 2 | 2600 ± 520 | 77000 ± 600 | -2.33 ± 0.48 |
| Metric 2 | 10 ± 4 | 4700 ± 1500 | 75000 ± 1500 | -2.71 ± 0.66 |
| Metric 3 | 10 ± 4 | 5000 ± 1600 | 74000 ± 1600 | -2.71 ± 0.68 |

Table S4: Mean first passage times (MFTPs) between the unbound (1) and the bound (2) state as estimated from the Bayesian MSM. MFTPs are measured in ns. The kinetic reaction rates of the three different force fields for the three metrics, $k_{on}$ and $k_{off}$ for the bound and the unbound states respectively. $k_{off}$ is reported in $\mu s^{-1}$ and $k_{on}$ in $\mu s^{-1}$ $M^{-1}$.

| SOMD/ff14SB/GAFF2 | | | | |
|---|---|---|---|---|
| Metrics | $MFTP_{1->2}$ | $MFTP_{2->1}$ | $k_{on}$ | $k_{off}$ |
| Metric 1 | 14.28 ± 0.04 | 15.03 ± 0.04 | 317.04 ± 0.83 | 7.00 ± 0.02 |
| Metric 2 | 14.76 ± 0.04 | 15.88 ± 0.04 | 300.08 ± 0.79 | 6.78 ± 0.02 |
| Metric 3 | 28.80 ± 0.20 | 9.30 ± 0.15 | 512.29 ± 3.65 | 3.47 ± 0.05 |
| GROMACS/Charmm36m/Charmm | | | | |
| Metric 1 | 11.32 ± 0.02 | 12.62 ± 0.03 | 373.69 ± 0.79 | 8.84 ± 0.01 |
| Metric 2 | 13.84 ± 0.03 | 13.95 ± 0.03 | 340.70 ± 0.67 | 7.17 ± 0.01 |
| Metric 3 | 7.37 ± 0.04 | 22.25 ± 0.10 | 212.02 ± 0.10 | 13.56 ± 0.07 |
| GROMACS/Amberidp/GAFF2 | | | | |
| Metric 1 | 13.10 ± 0.02 | 28.93 ± 0.08 | 165.39 ± 0.39 | 7.63 ± 0.02 |
| Metric 2 | 15.84 ± 0.04 | 35.11 ± 0.12 | 136.23 ± 0.33 | 6.31 ± 0.02 |
| Metric 3 | 16.59 ± 0.08 | 14.98 ± 0.13 | 319.36 ± 2.73 | 6.03 ± 0.05 |

### SUPPLEMENTARY FIGURES

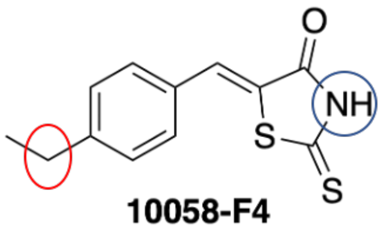


10058-F4

Figure S2: Structure of 10058-F4. The two atoms of the molecule that are circled were chosen to measure distances between the ligand and the alpha carbons of the c-Myc peptide. These distances were used as molecular features for the MSM models.

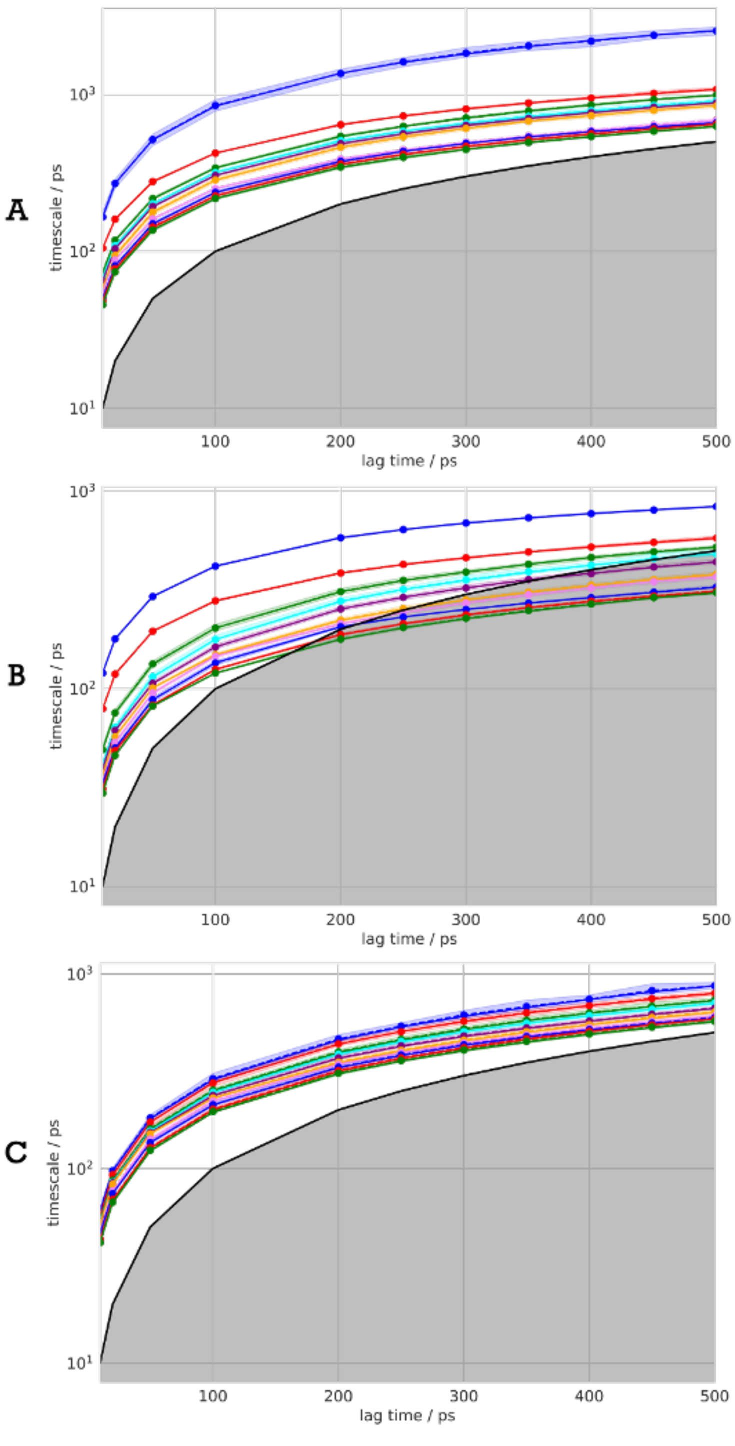


Figure S3: Implied time scales plots of the first metric for the three force fields, **A** AmberIDP, **B** Charmm36m, **C** FF14SB. Different colors indicate the slowest processes of the system during the MD simulations.

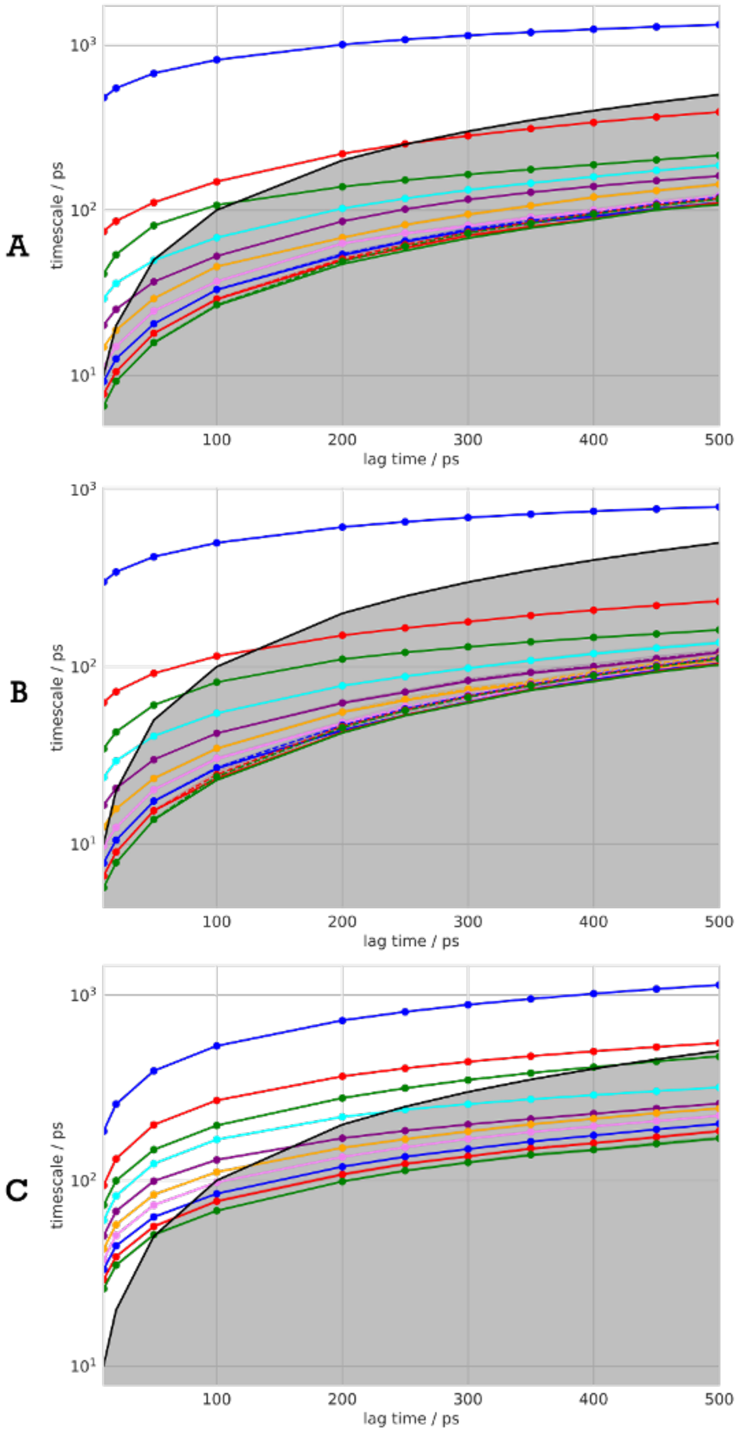


Figure S4: Implied time scales plots of the second metric for the three force fields, **A** AmberIDP, **B** Charmm36m, **C** FF14SB. Different colors indicate the slowest processes of the system during the MD simulations.

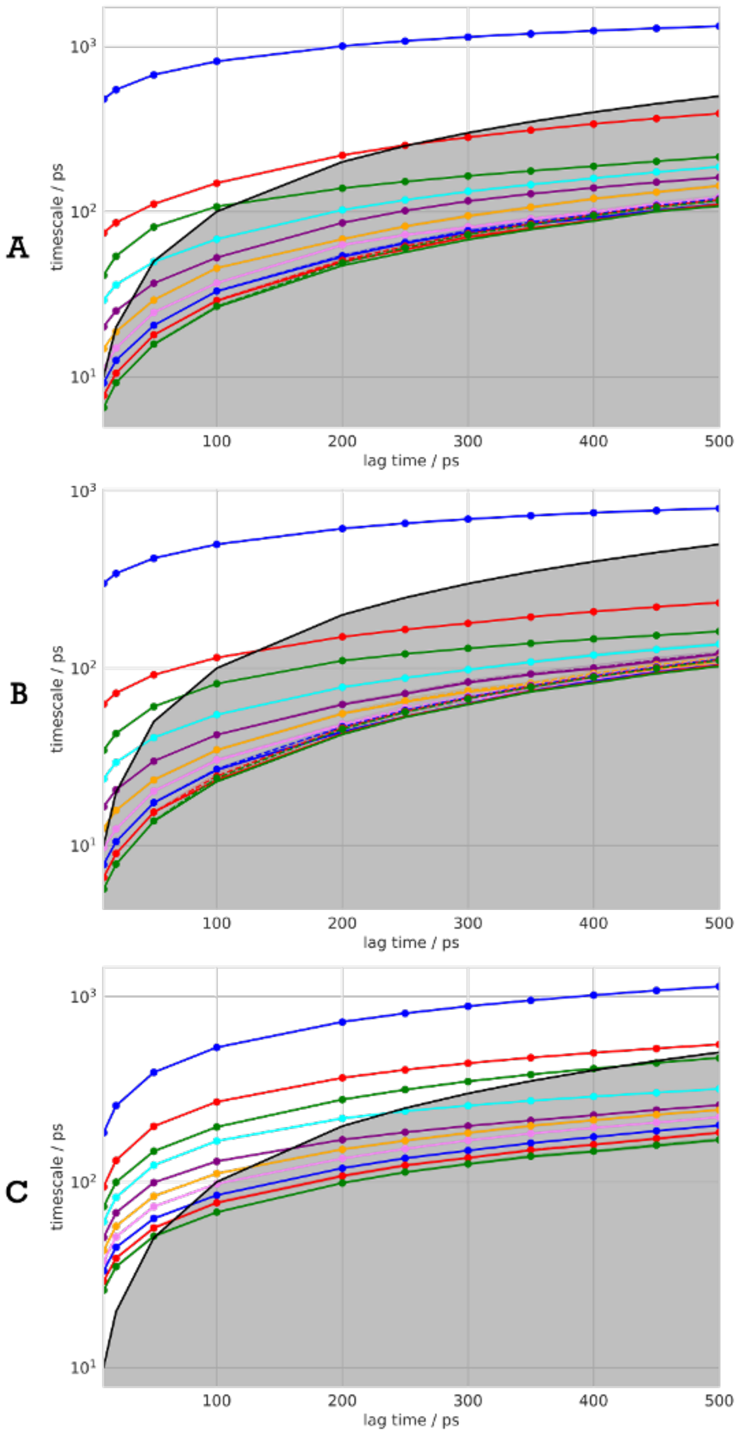


Figure S5: Implied time scales plots of the third metric for the three force fields, **A** AmberIDP, **B** Charmm36m, **C** FF14SB.

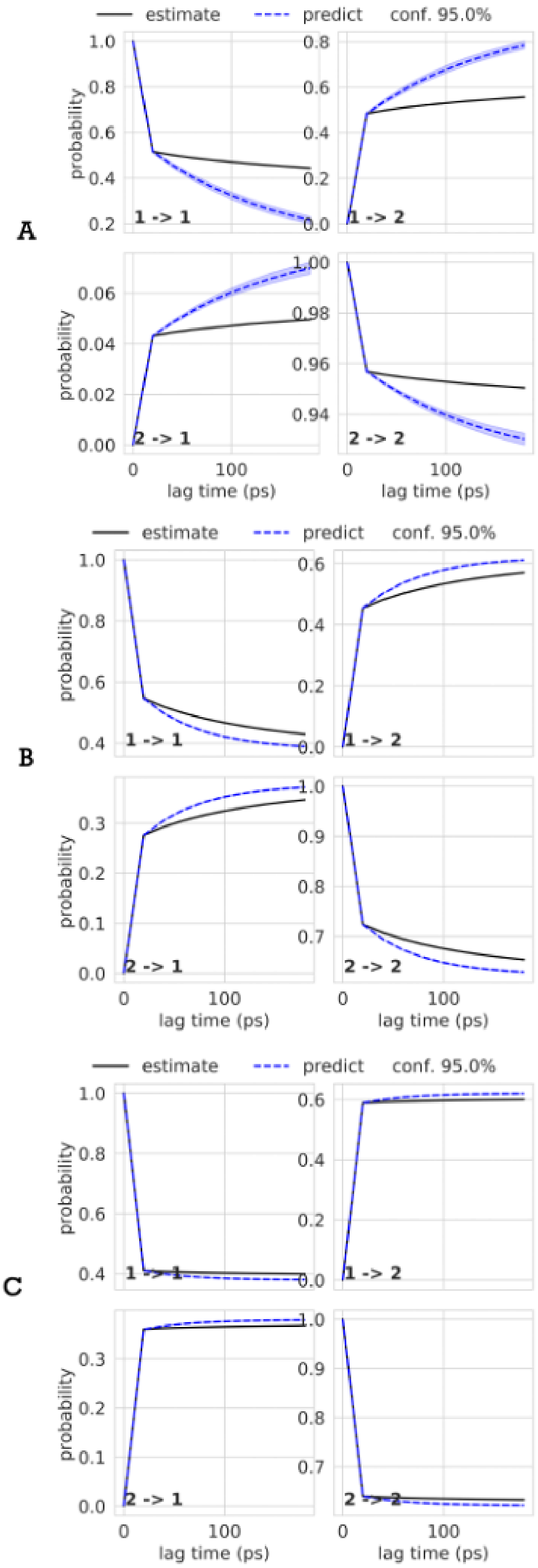


Figure S6: Chapman-Kolmogorov test plots of the first metric used for the three force fields, **A** AmberIDP, **B** Charmm36m, **C** FF14SB.

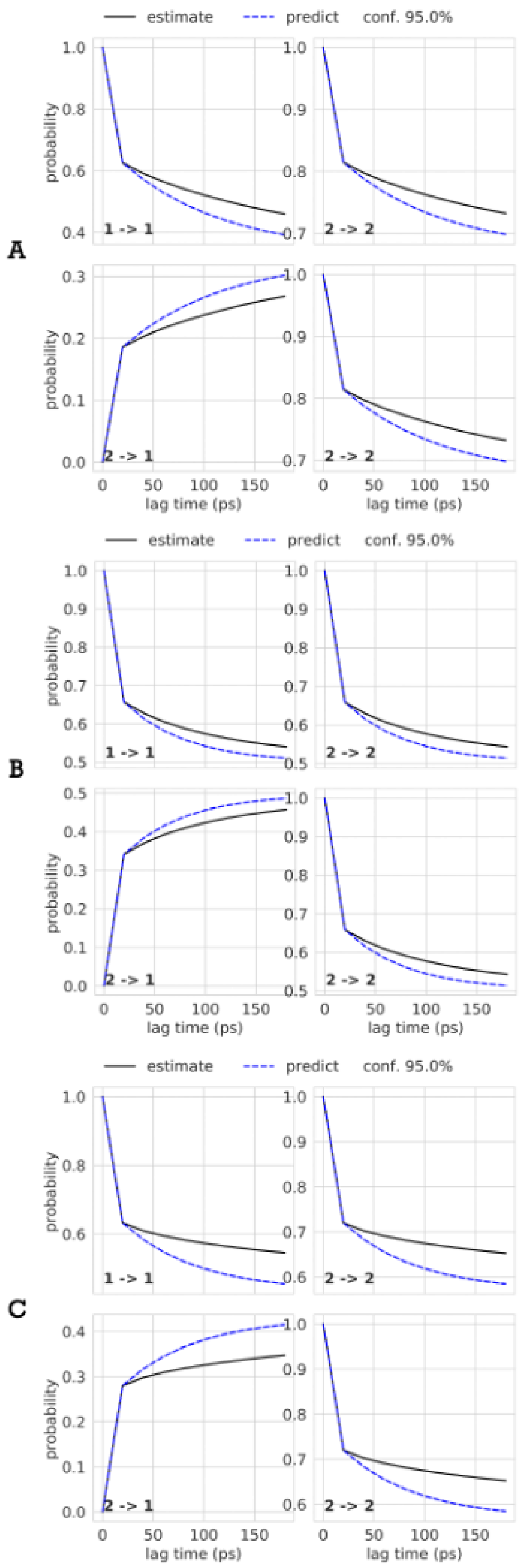


Figure S7: Chapman-Kolmogorov test plots of the second metric used for the three force fields, **A** AmberIDP, **B** Charmm36m, **C** FF14SB.

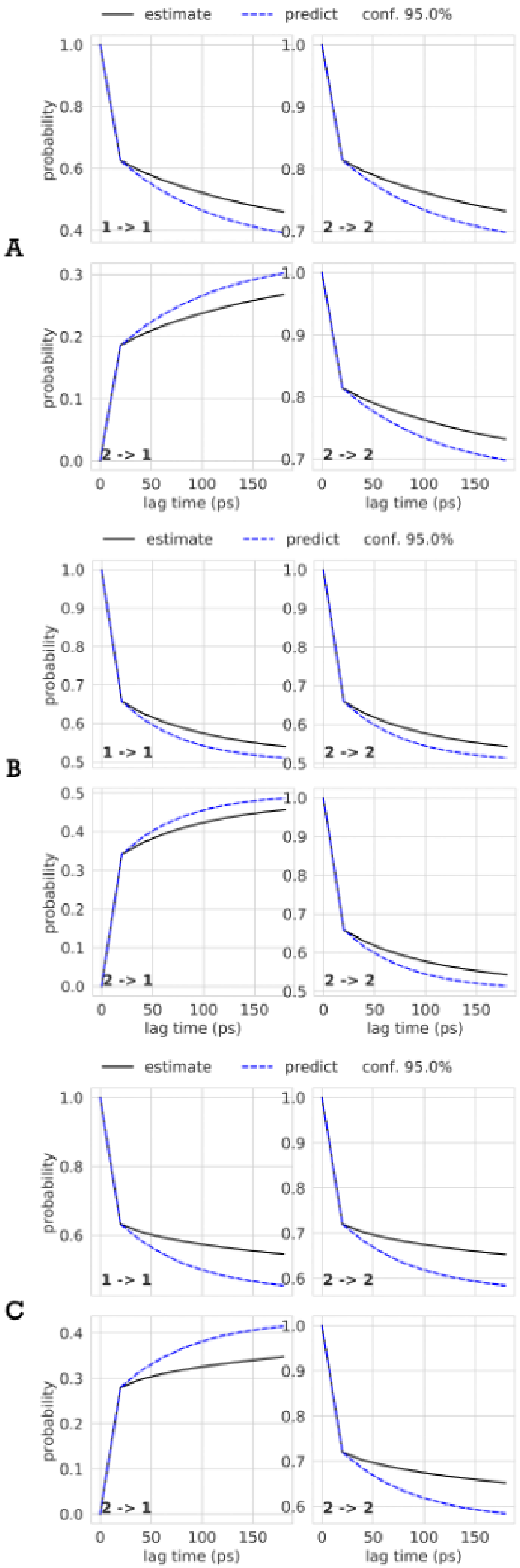


Figure S8: Chapman-Kolmogorov test plots of the third metric used for the three force fields, **A** AmberIDP, **B** Charmm36m, **C** FF14SB.

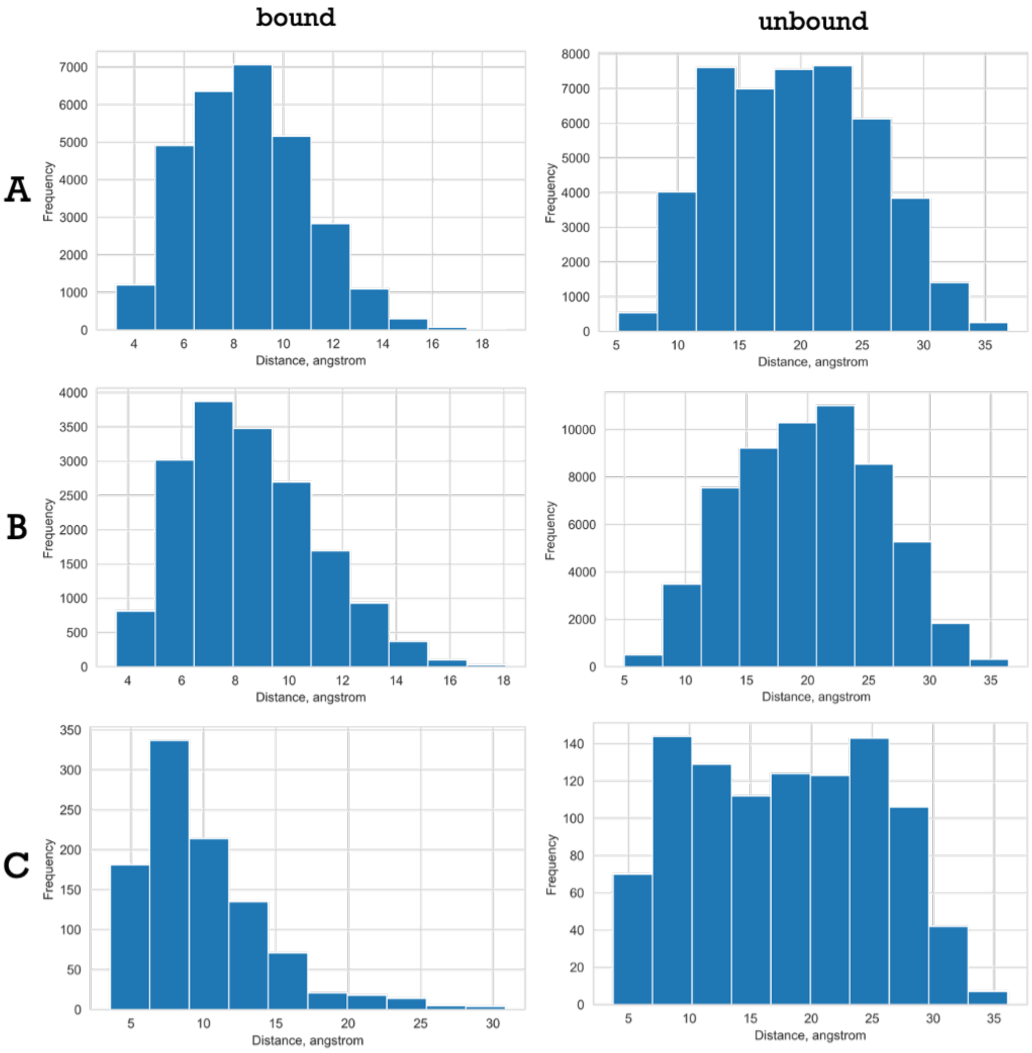


Figure S9: Probability distribution of distances of the bound and unbound states of the first metric applied for the three force fields, **A** AmberIDP, **B** Charmm36m, **C** FF14SB.

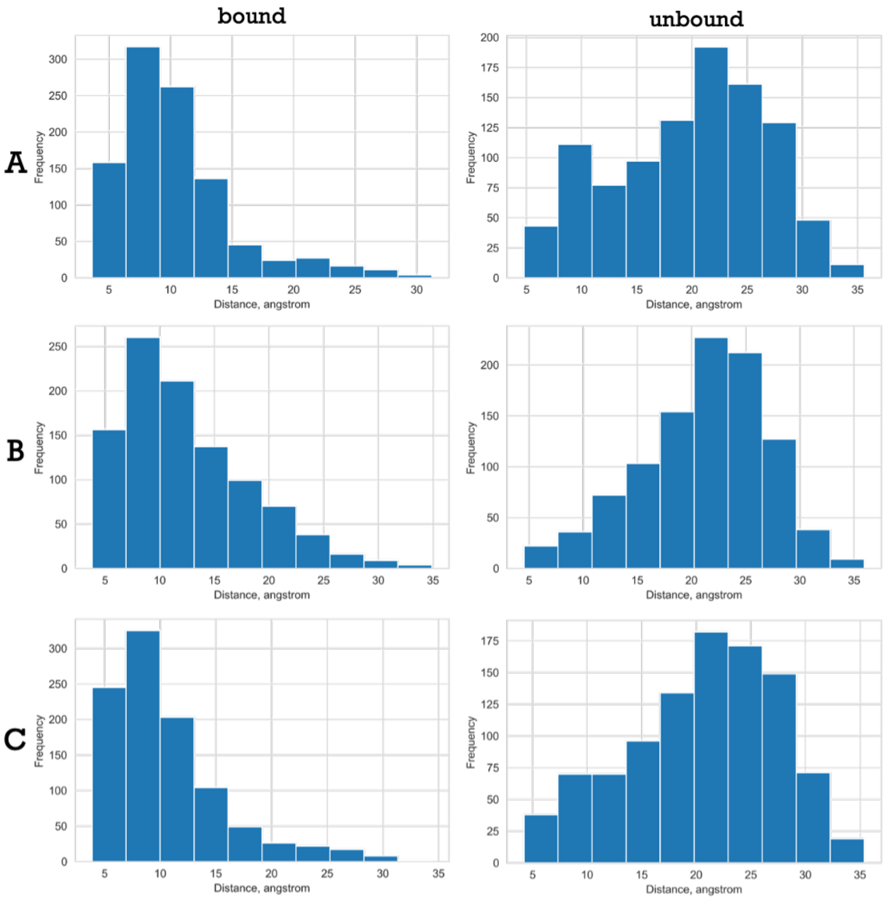


Figure S10: Probability distribution of distances of the bound and unbound states of the second metric applied for the three force fields, **A** AmberIDP, **B** Charmm36m, **C** FF14SB.

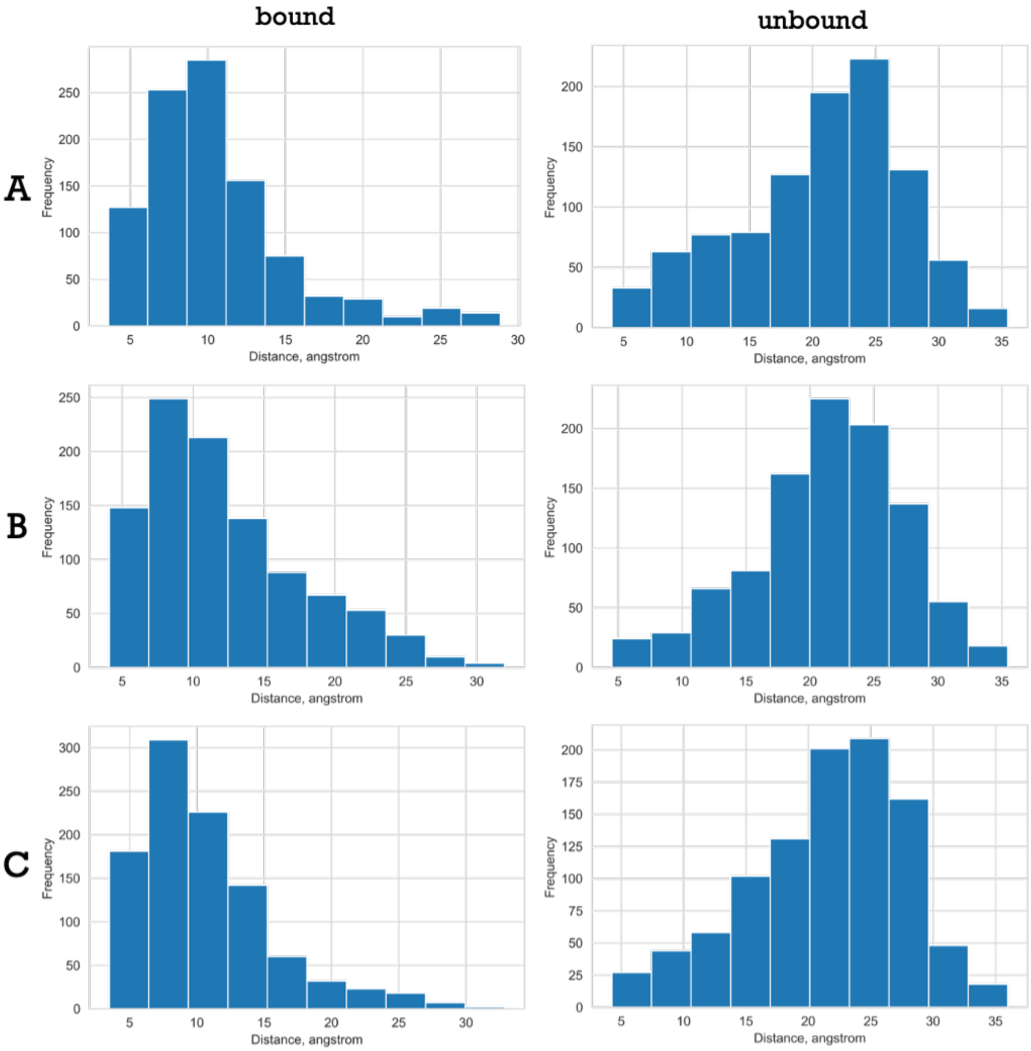


Figure S11: Probability distribution of distances of the bound and unbound states of the third metric applied for the three force fields, **A** AmberIDP, **B** Charmm36m, **C** FF14SB.

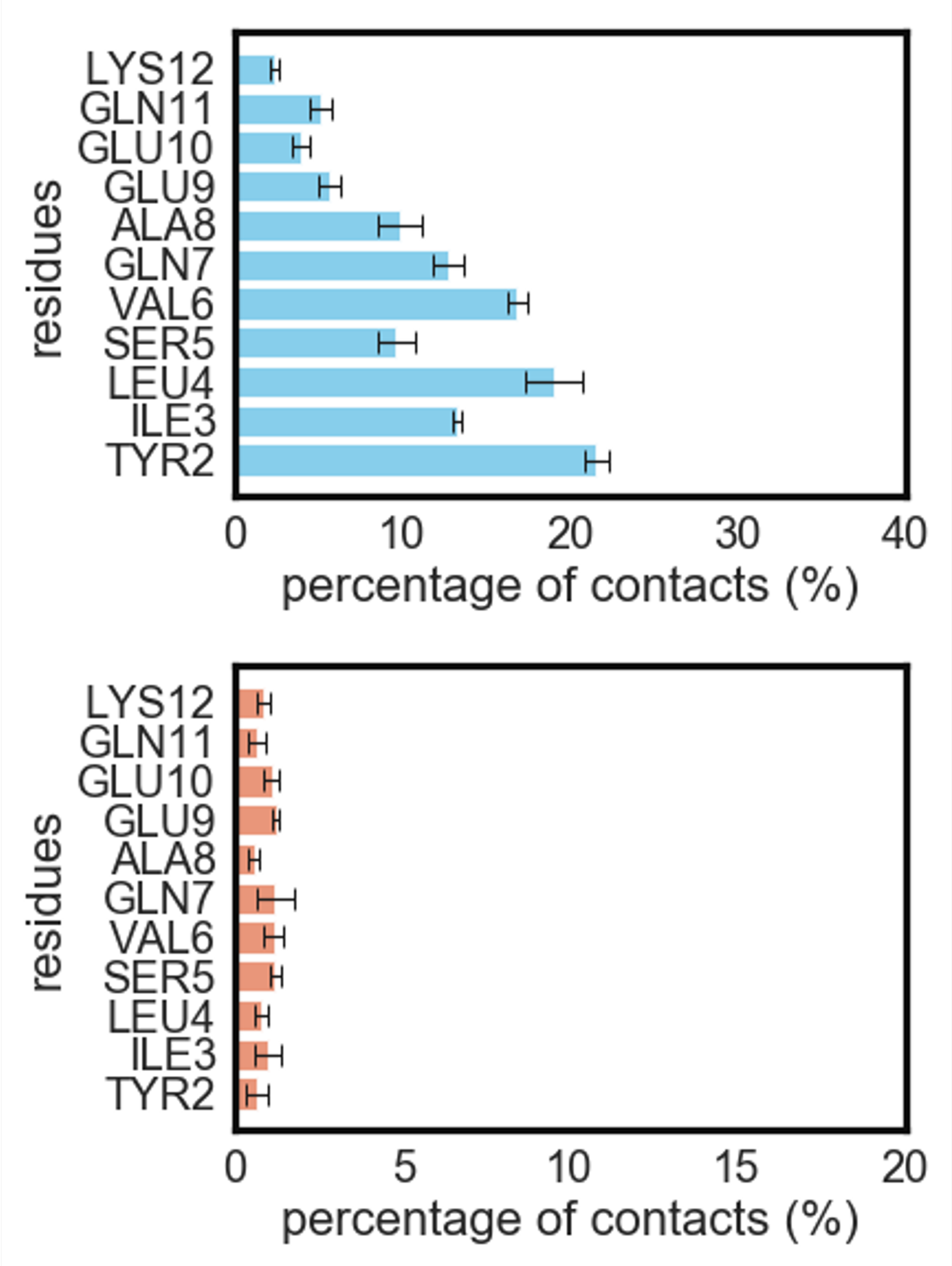


Figure S12: A) Mean percentage of hydrophobic contacts formed during the four trajectories, each containing 1000 snapshots and B) Mean percentage of hydrogen bonds formed during the four trajectories, each containing 1000 snapshots for Charmm force-field.

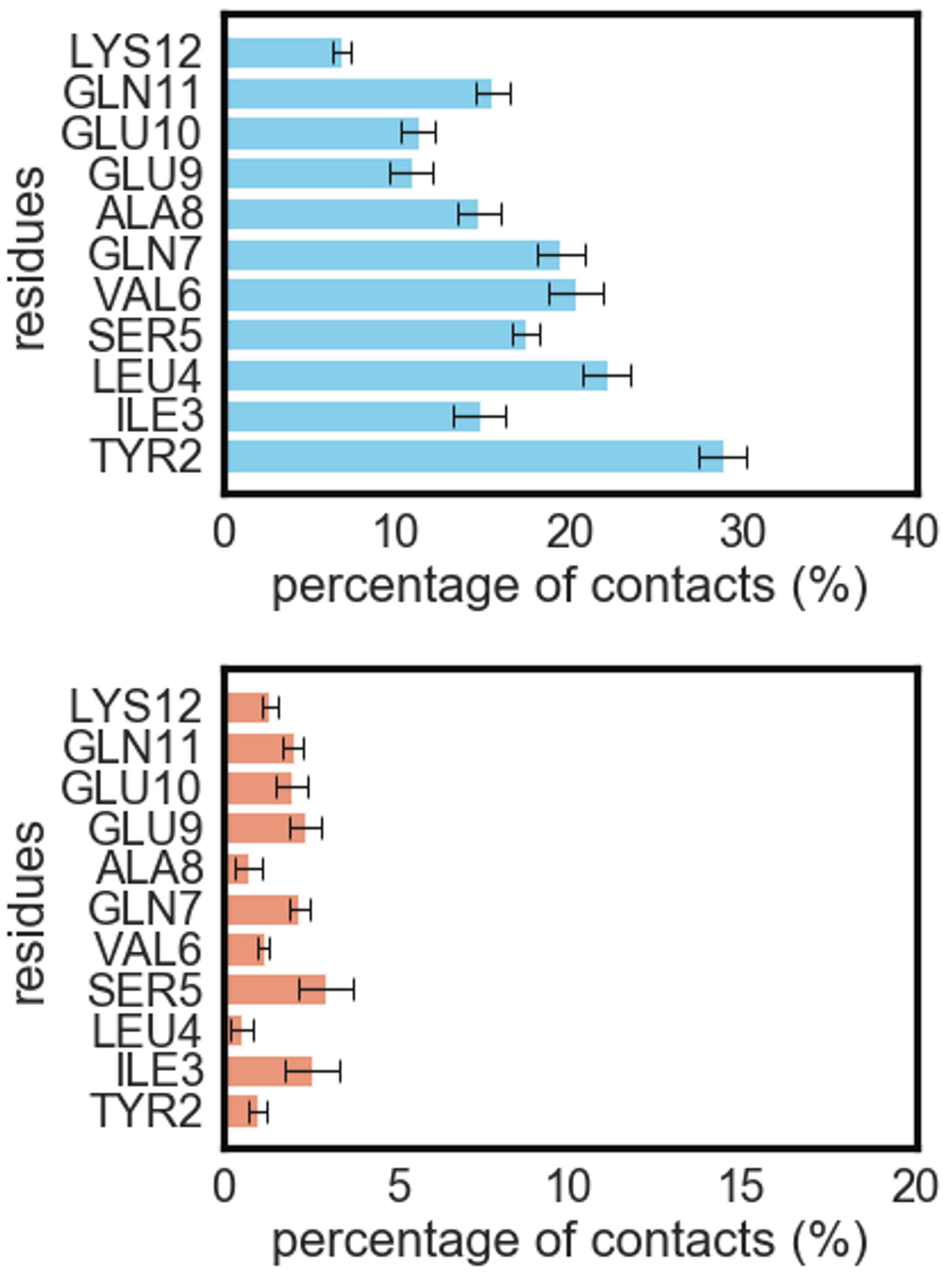


Figure S13: A) Mean percentage of hydrophobic contacts formed during the four trajectories, each containing 1000 snapshots and B) Mean percentage of hydrogen bonds formed during the four trajectories, each containing 1000 snapshots for ff14SB force-field.
